# Metabolic rewiring of the probiotic bacterium *Lacticaseibacillus rhamnosus* GG contributes to cell-wall remodeling and antimicrobials production

**DOI:** 10.1101/2023.01.03.522566

**Authors:** Ronit Suissa, Tsviya Olender, Sergey Malitsky, Ofra Golani, Sondra Turjeman, Omry Koren, Michael M. Meijler, Ilana Kolodkin-Gal

**Affiliations:** Department of Chemistry, Ben-Gurion University of the Negev, Be’er Sheva, Israel; Department of Molecular Genetics, Weizmann Institute of Science, Rehovot, Israel; Life Science Core Facilities, Weizmann Institute of Science, Rehovot, Israel; Azrieli Faculty of Medicine, Bar-Ilan University, Safed, Israel; Department of Plant Pathology and Microbiology, Faculty of Agriculture, Food and Environment, The Hebrew University of Jerusalem, Rehovot, Israel

## Abstract

*Lacticaseibacillus rhamnosus GG* (LGG) is a Gram-positive beneficial bacterium that resides in the human intestinal tract and belongs to the family of lactic acid bacteria (LAB). This bacterium is a widely used probiotic and was suggested to provide numerous benefits for human health. However, as in most LAB strains, the molecular mechanisms that mediate the competitiveness of probiotics under different diets remain unknown. Fermentation is a fundamental process in LAB, allowing the oxidation of simple carbohydrates (e.g., glucose, mannose) for energy production under conditions of oxygen limitation, as in the human gut. Our results indicate that fermentation reshapes the metabolome, volatilome, and proteome architecture in LGG. Furthermore, fermentation alters cell envelope remodeling and peptidoglycan biosynthesis, which leads to altered cell wall thickness, aggregation properties, and cell wall composition. In addition, fermentable sugars induced secretion of known and novel metabolites and proteins targeting the enteric pathogens *Enterococcus faecalis* and *Salmonella Enterica serovar Typhimurium*. Overall, our results link the common metabolic regulation of cell wall remodeling, aggregation to host tissues, biofilm formation in probiotic strains, and connect the production of antimicrobial effectors with metabolome reprogramming. These findings provide novel insights into the role of nutrition in the establishment of LGG in the gastrointestinal tract.

## Introduction

Probiotic strains are consumed either as fresh fermentation products or as dried bacterial supplements with Lactobacillaceae and Bifidobacteria, being the two widely used probiotic genera (1, 2). The main strains currently used for probiotic formulation were originally isolated from fermented products or humans (3–5). Lactobacillaceae are Gram-positive rods, facultative anaerobes, and belong to the lactic acid bacteria (LAB) group, as lactic acid is their main end-product of carbohydrate metabolism (6, 7). They are naturally found in the gastrointestinal tract (GIT) of humans and animals, as well as in the urogenital tract of females (8)and are widely used probiotic bacteria, represented in most fermented products and supplements. Probiotic performance in the gut has been shown to be dependent on nutrient composition and availability(9).

LAB are considered efficient fermenters, proficient in producing energy under anaerobic conditions or when oxygen is limited. The fermentation process involves the oxidation of carbohydrates to generate a range of products, including organic acids, alcohol, and carbon dioxide (10). The response of all LAB strains to fermentation is considered uniform and primarily depends on the capacity to utilize glucose and its products (2). However, we recently found that the response of *Lacticaseibacillus rhamnosus GG* (LGG) to fermentable sugars is strain-specific and includes regulatory responses that are independent of cell growth (11). Glucose utilization has a general role in shaping the colony morphology under static conditions, as all species exhibit morphological colony changes upon glucose treatment(11). Interestingly, buffering of the media restores glucose-free morphology of all strains, indicating that acid stress is involved in biofilm formation throughout the Firmicutes phylum directly or indirectly (11). Glucose-induced change in colony morphology occurred independently of cell death and fermentation-associated pH drop. Therefore, they may be reflective of a complex adaptation and biofilm regulation, rather than cell density. In addition, pH independent changes occurred in the aggregation and adhesion properties of the LAB strains tested with glucose. These results indicate that specific responses to carbohydrates as mediators of cellular adhesive properties could be attributed to metabolic and proteomic rewiring. Here we chose the proficient probiotic bacterium LGG as our lens, to unravel the molecular mechanisms that act downstream to glucose metabolism and contribute to its altered fitness upon glucose utilization.

## Results

### The effect of fermentable sugars on proteome architecture

We previously found that fermentable sugars specifically effect the probiotic bacterium LGG, functioning as distinct regulators of growth, aggregation and adhesion(11). The application of glucose, galactose and mannose, which can be fermented by LGG (12) induced growth compared to the tryptic soy broth (TSB) control medium. Here, we extended the repertoire of consumed carbohydrates in LGG. The application of starch increased the growth of LGG to a lesser extent compared with glucose, galactose, and mannose. Starch is composed of glucose units joined by the glyosidic bond, and this result indicates that LGG can only partially degrade starch into glucose. However, non-fermentable sugars such as sucrose, xylose and raffinose did not induce growth of LGG in shaking cultures (Figure 1A). The effects of fermentation on bacterial growth and pH are well established. However, the metabolic status of bacteria in the GI can also affect gene expression, and therefore the protein architecture. An unbiased proteomics analysis was performed to test the global response of LGG to fermentation, and to identify pathways that are changed following fermentation. To the best of our knowledge, this is the first proteomic evaluation of pathways acting downstream to fermentation on LGG. Principal component analysis (PCA) of the proteomic data demonstrated that the protein profiles of LGG in fermentation (application of glucose and mannose), are clustered closely together but both partition from the proteome of non-fermenting cultures (application of raffinose or TSB control) with the first component covering 40% of the data variance (Figure 1B). This result is also reflected in the heat map, where two major clusters separating between fermentation and non-fermentation conditions can be seen (Figure 1C). Therefore, fermentable sugars trigger profound changes in the physiology of LGG. We determined that these changes are triggered by fermentation and not specifically by glucose as a carbon source, as both mannose and glucose are converted into fructose-6-phosphate and proceed to the glycolysis pathway (10).

**Figure 1.**
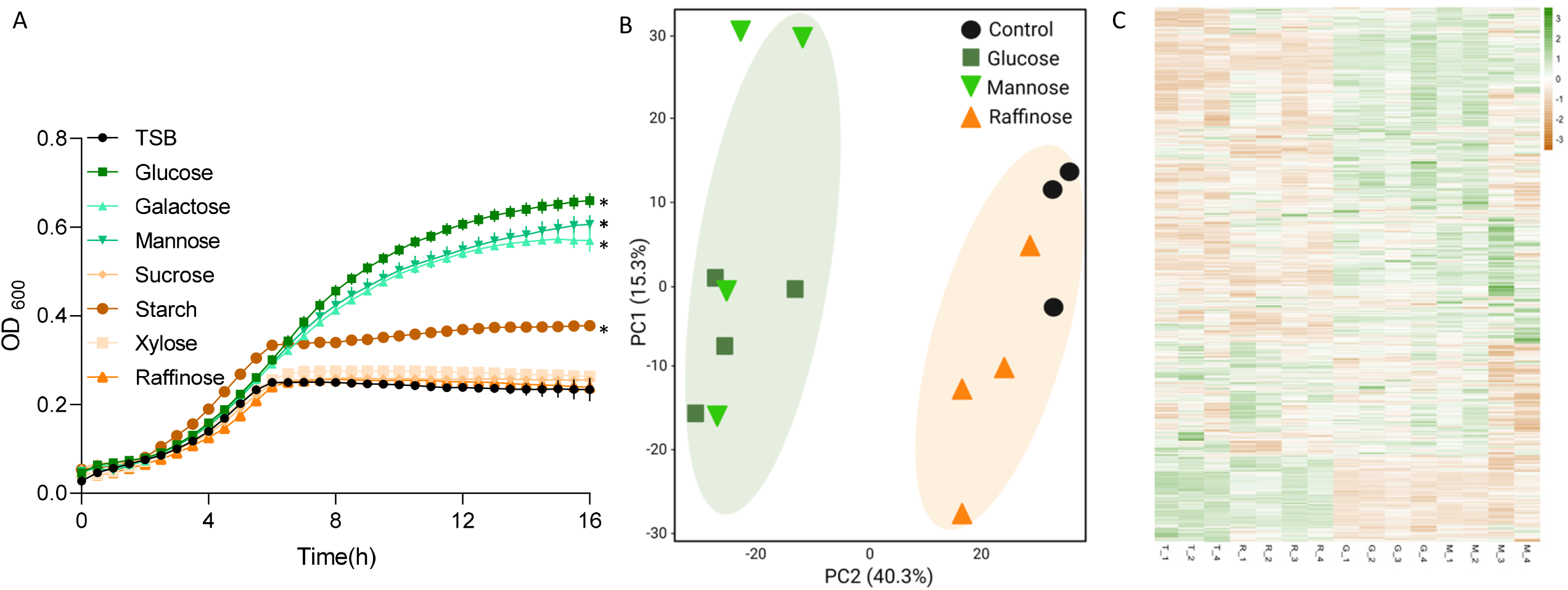
**(A)** Growth curves of LGG in 96 well plates at 37°C with shaking in TSB medium or TSB medium supplemented with 1% different sugars. Statistical analysis was performed using one-way ANOVA with Tukey’s post-hoc test vs. TSB. p < 0.05 was considered statistically significant. **(B)** Principal component analysis (PCA) plot of proteomics analysis of LGG grown in liquid TSB medium [T1-T4] or TSB medium supplemented with glucose (1% W/V) [G1-G4], mannose (1% W/V) [M1-M4] or raffinose (1% W/V) [R1-R4]. **(C)** Proteins heat map which are based on fold differences in intensity of all proteins identified in proteomics analysis.

### Metabolome rewiring is associated with proteome remodeling

To link between genes and responses of biological systems to the environmental variations we performed untargeted metabolomics on LGG grown with and without glucose and mannose. As judged by PCA, LGG cultured with fermentable sugars clustered separately from the control media and raffinose supplemented media, with the first component, covering 44.5% of the data variance (Figure S1). These results correlate with our proteomics results (Figure 1B). The MetaCyc metabolic pathway toolkit (13) indicated that metabolites from the pyruvate fermentation pathways changed significantly in response to the carbohydrate source (Figure 2A-C). Pyruvate abundance in LGG cultured with glucose and mannose was significantly higher than un-supplemented media. However, pyruvate also increased with raffinose, suggesting that under our experimental conditions, pyruvate is not directly affected by fermentation (Figure 2D). Both lactic acid and acetyl-CoA were significantly induced (compared with TSB control) with glucose and mannose, but not raffinose (Figure 2E,F). These results indicate that lactic acid and acetyl-CoA may be the cellular cues that are translated into the specific changes in the LGG proteome.

**Figure 2.**
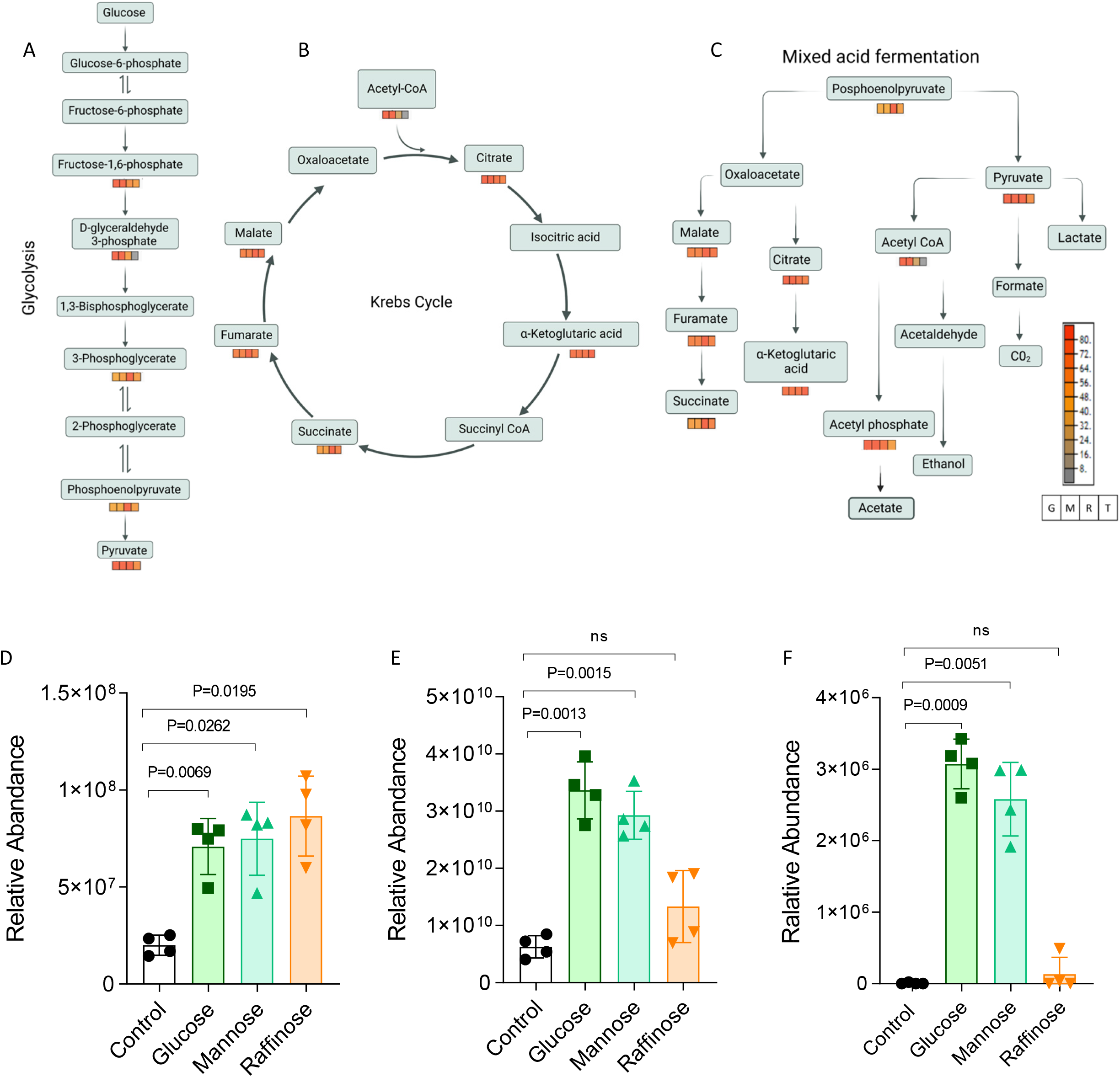
LGG metabolites profile is affected by fermentation. **(A-C)** MetaCyc metabolic pathway of fermentation based on metabolomics analysis from LGG grown in TBS medium or TSB medium with glucose (1% W/V), mannose (1% W/V) or raffinose (1% W/V). Relative abundance of **(D)** Pyruvate **(E)** Lactic acid **(F)** Acetyl-CoA. The images represent 4 independent repetitions. Statistical analysis was performed using Brown-Forsythe and Welch’s ANOVA with Dunnett’s T3 multiple comparisons test. p < 0.05 was considered statistically significant.

Changes in intracellular metabolome also effect the air borne metabolites that mediate interspecies interactions from a far, also termed volatiles. To test whether extracellular volatiles are affected by carbohydrate consumption and how, we systematically analyzed the LGG volatilome, considering the effects of fermentation on this process. We grew LGG in liquid TSB or TSB applied with glucose and sampled the Volatile compounds (VCs) in the headspace above the cultures using a GC-TOF-MS. Consistent with the metabolic rewiring of the metabolome and proteome, fermentation altered the VCs profile and overall induced the production of organic volatiles (Figure 3A). In the TSB medium, background from the media was evident for multiple volatiles, complicating the analysis. However, 1-Butanol (Figure 3C) and 3-Methylbutanal (Figure 3D) were reduced during fermentation, while Isoamyl alcohol (Figure 3E) was increased by fermentation and pyrazine was produced in detectable amount with and without the fermentable sugar, glucose (Figure 3F). To reduce background, we measured the volatilome again in a defined medium (Figure 3B) with and without glucose as the added carbohydrate. 1-Butanol (Figure 3G), Diacetyl (Figure 3H) and 3-Methylbutanal (Figure 3I) were downregulated during fermentation. Acetic acid ethenyl ester was induced during fermentation (Figure 3J), coincident with the increase in cytoplasmic acetic acid (Figure 2) and pyrazine was produced in significant amount in both the absence and presence of glucose (Figure 3K). These results indicate that the volatilome of LGG is not a direct readout of the cellular metabolome, and has a more complicated response to the fermentable sugar, glucose. Nevertheless, the significant differences in the emitted volatiles in the presence of fermentable sugar (Figure 3A and B) confirm that fermentation has a cardinal role in shaping the exudated metabolome of LGG.

**Figure 3.**
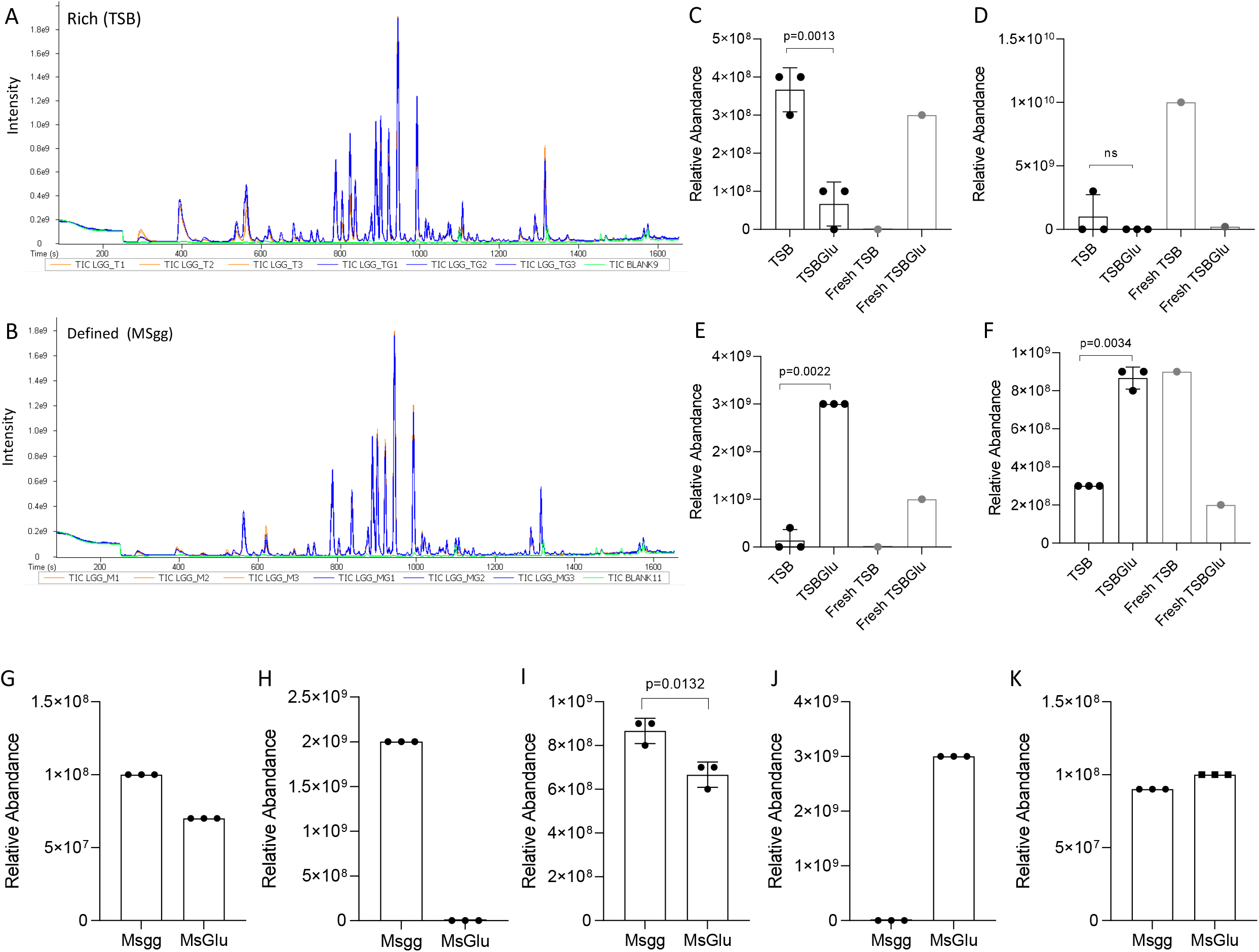
Discovery of fermentation-dependent volatiles. **(A)** Chromatographic profile obtained by GC-MS of volatile compounds from LGG grown with or without glucose in TSB or **(B)** defined medium Msgg. Relative abundance of compound identified in rich medium TSB; **(C)**1-Butanol **(D)** 3-Methylbutanal **(E)** Isoamyl alcohol **(F)** Pyrazine. Gray bars represent the background (fresh medium). Relative abundance of compound identified in defined medium Msgg; **(G)** 1-Butanol, **(H)** Diacetyl **(I)** 3-Methylbutanal **(J)** Acetic acid ethenyl ester **(K)** Pyrazine. Statistical analysis was performed using unpaired t-test with Welch’s correction. In G, H, J and K the three independent repetitions measurements gave identical values.

### Specific changes in cell wall biosynthesis are induced by fermentable sugar

We demonstrated that the intracellular and extracellular metabolome is rewired during fermentation and an enhanced intracellular accumulation of acidic products (Figure 2). In addition, we detected an increased emission of a volatile targeting the cell envelope (Acetic acid ethenyl ester) (Figure 3). Thereby to adapt to fermentable sugars, the bacteria are expected to increase their overall tolerance to cell envelope damage. Globally, about 195 out of 1800 proteins where significantly more abundant when the fermentable sugars glucose and raffinose were applied (Figure 4A). Therefore, we employed the PSORT analysis and WEB-based Gene Set Enrichment Analysis Toolkit (14), to explore potential microbial adaptations to the envelope stress exerted by fermentation products. Our analysis revealed that the cell wall biosynthesis and turnover components are induced by fermentation (Figure 4B, C). The Gram-positive bacterial cell wall is composed of layers of <u>peptidoglycan</u>, a polymer made from <u>polysaccharide</u> chains cross-linked by <u>peptides</u> containing D-<u>amino acids</u>, and layers of teichoic acid (15). The categories of cell wall proteins that were altered include penicillin binding proteins, which are generally involved in peptidoglycan biosynthesis(16).

**Figure 4.**
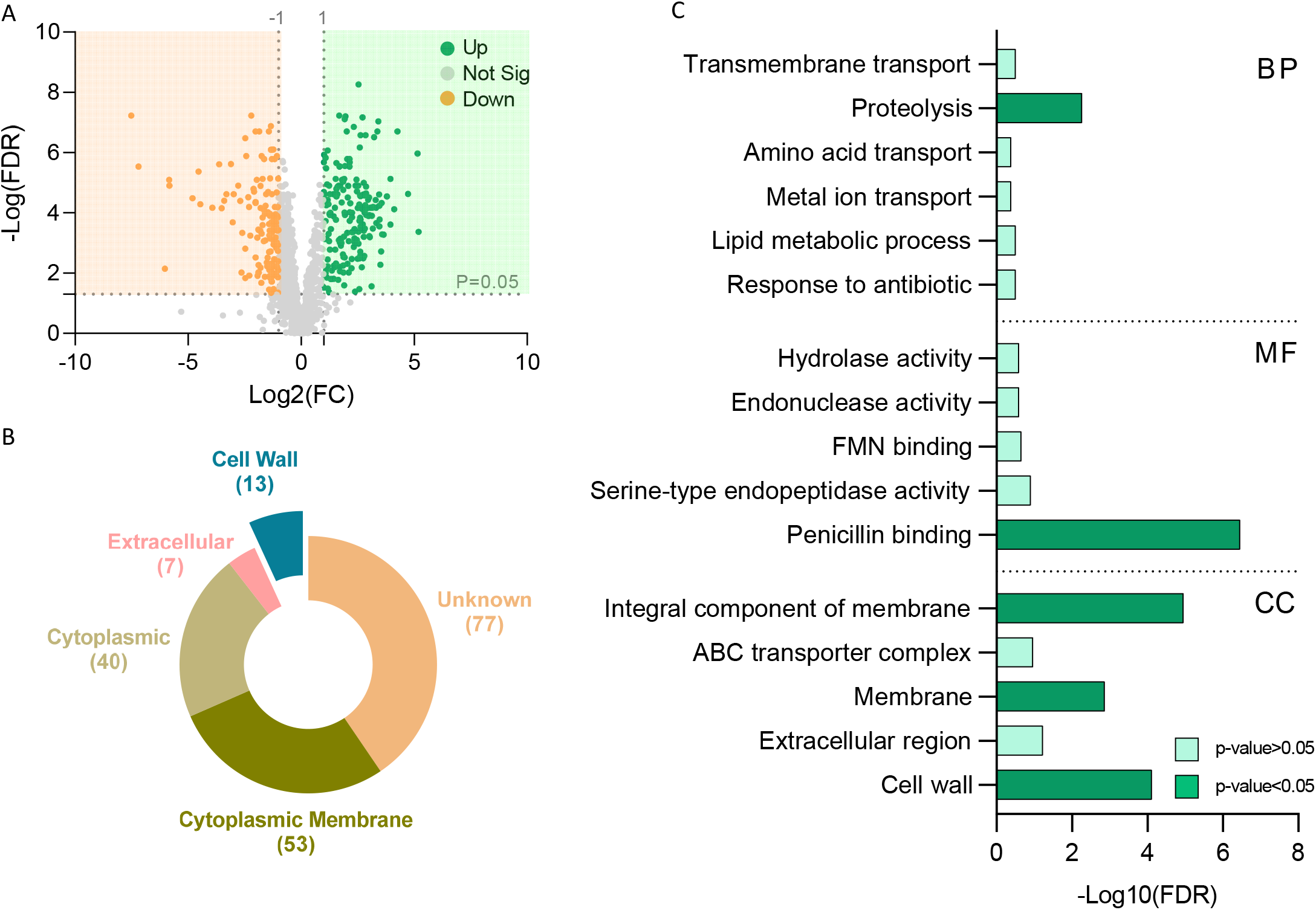
**(A)** Volcano plot representing the log2 fold-change (fermentation/non-fermentation) against -log10 statistical p value with FDR correction for proteins differentially expressed between glucose and mannose vs. control and raffinose. Gray dots represent unchanged proteins, green and orange dots represent the upregulated and downregulated proteins respectively; the horizontal dashed line indicates FDR = 0.05 and vertical dashed line indicates the log2FC. (B) Pie chart represent PSORT analysis and **(C)** Bar chart shows Over Representation Analysis (ORA-WebGestalt) of upregulated proteins during fermentation.

To further evaluate the response of the cell wall to fermentation, we manually screened for cell-wall proteins that may affect the competitiveness of LGG in the GI and are affected by fermentation. We found that a class A Sortase (Figure S2) induced by fermentation. This membrane-associated cysteine transferase belongs a family of proteins that facilitate the bacterial evasion of the host’s immune system (17) and in immunomodulation, adhesion to epithelial cells or antibacterial activity against gut pathogens (18). To confirm that indeed the cell wall is remodeled during fermentation, we studied LGG stained with green, fluorescent membrane dye (FM4-64) (Axio microscope, Zeiss, Germany). We found that LGG cells grown in colonies in the absence of fermentable sugar are rod shaped and clearly separated (Figure 5A). In contrast, colony cells grown with fermentable sugar (Figure 5A) lost the elongated rod shape. To get a higher resolution of the cell wall morphology changes of cells grown with and without a fermentable sugar, we looked at the cells under transmission electron microscopy (TEM) which allows us to resolve cellular structures by performing a cross-section of the bacterial cells. The cells grown without glucose had a rod shape with smooth edges. In contrast, cells grown on glucose have a rounder atypical shape, consistent with their appearance in the light microscope. The edges are not smooth but rather laced (Figure 5B). We suggest that the changes that we observed in the proteome may explain the underlying mechanism for the observed changes in the cells. A quantification of cell wall thickness cell from TEM images (Figure 5C) confirmed this adaptation, consistent with enhanced tolerance to antimicrobials(19). In addition, image stream flow cytometry was also indicating significant alterations in the cellular envelope (Figure 5D).

**Figure 5.**
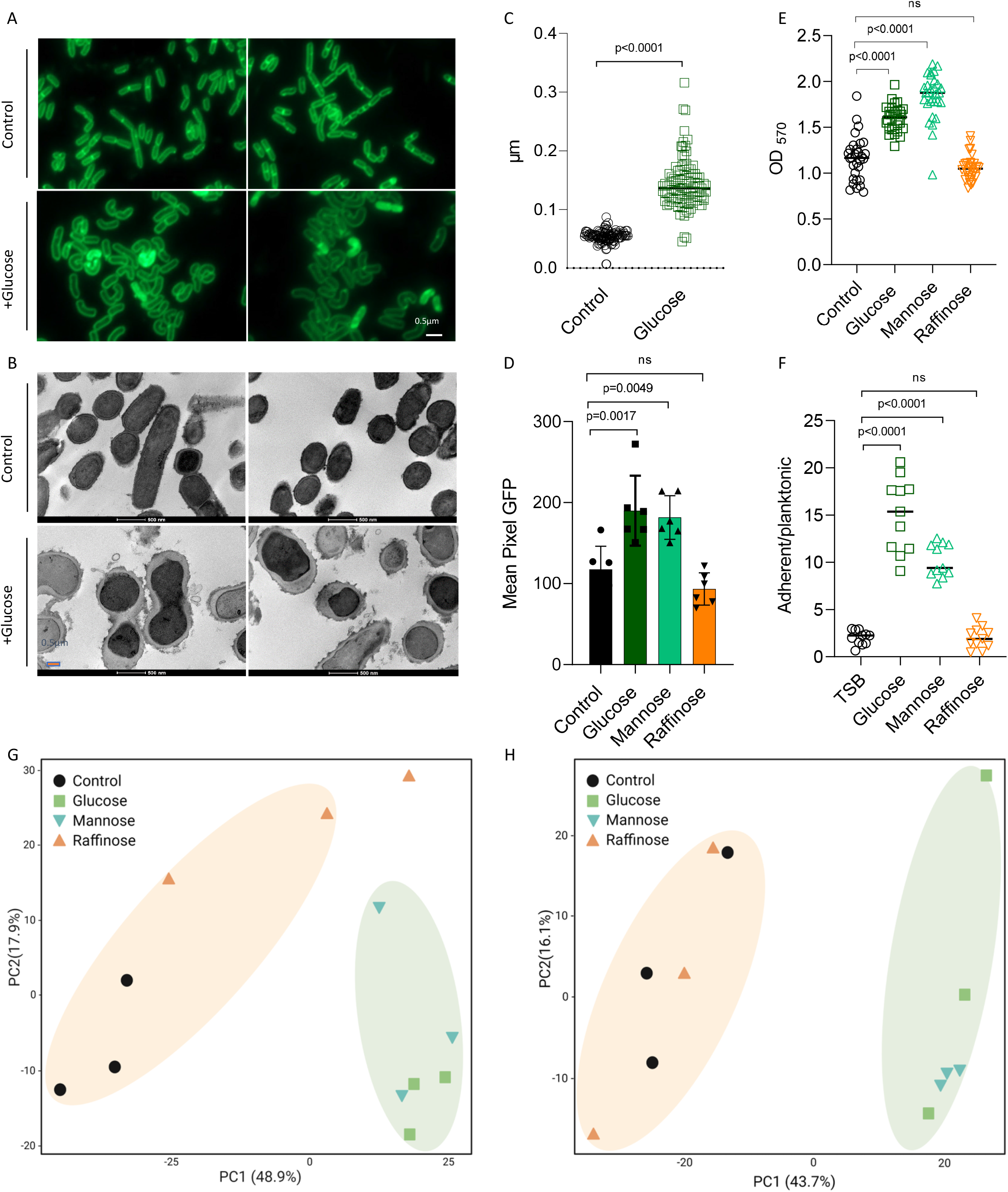
**(A)** Fluorescence microscope images of cells from LGG colonies that were grown on solid TSB (control), TSB supplemented with glucose (1% W/V). Cells were stained using Green-membrane stain FM™ 1–43FX. Biofilms were grown at 37° C in CO2 enriched environment. Cells were imaged at 72 h post inoculation. The images represent 3 independent experiments, from each repeat at least 10 fields examined. Scale bar = 2 μm. **(B)** Transmission electron microscopy (TEM) images of LGG colony grown in solid TSB medium or TSB medium (control) supplemented with glucose (1% W/V). Scale bar=0.5 μm. **(C)** Quantification of 100 cells cell wall thickness from TEM images using Fiji-ImageJ. **(D)** Imaging Flow cytometry analysis of the mean pixel intensity of LGG colony that were grown on solid TSB (control), TSB supplemented with glucose (1% W/V), mannose (1% W/V) or raffinose (1% W/V). Cells were labeled using WGA-FITC. Biofilms were grown at 37° C in CO_2_ enriched environment. Data were collected after 72h, and 100,000 cells were counted. Graphs represent mean ± SD from 2 independent experiments (n = 6). **(E)** LGG cells were diluted 1:100 into a fresh TSB medium or TSB supplemented with glucose (1% W/V), mannose (1% W/V) or raffinose (1% W/V). 200 μL of cultures were split into a 96-well polystyrene plate and further incubated at 37 °C. Crystal violet assay assessed the biofilm formation of LGG after 72h. Graph represent the mean ± SD from three biological repeats (*n* = 30). **(F)** LGG cells were diluted 1:100 into a fresh TSB medium or TSB supplemented with glucose (1% W/V), mannose (1% W/V) or raffinose (1% W/V). 1.5mL of cultures were split into a 12-polystyrene plate and further incubated at 37 °C for 72h. The upper growth media was removed and OD_600_ measured. The remaining biofilm biomass was diluted in fresh TSB and OD_600_ measured. Y axis represent the ratio of OD_600_ (planktonic/biofilm biomass). Graph represent the mean ± SD from three biological repeats (*n* = 12). All Statistical analysis was performed using Brown-Forsythe and Welch’s ANOVA with Dunnett’s T3 multiple comparisons test. p < 0.05 was considered statistically significant. **(G)** PCA analysis compering between treatments of biofilm-derived peptidoglycan and **(H)** planktonic-derived peptidoglycan. The images represent 3 independent repetitions.

In non-LAB bacteria, it was demonstrated that cell wall homeostasis is tightly linked to biofilm formation (20, 21) and is frequently co-regulated with the extracellular matrix genes (22–24). Therefore, we tested biofilm formation across the tested carbohydrates. Indeed, glucose and mannose but not raffinose were significantly inducing biofilm formation as judged by the biofilm biomass accumulating on the cell surface (Figure 5E), and the ratio of surface-associated and free-living cells (Figure 5F). Under these conditions the colony was flat and featureless. Bacteria positioned at the edges of the colony swarmed [collectively migrate over the agar surface], and, therefore, the edges of the colony were not symmetrical. The application of raffinose had little or no effect on colony morphology. However, the application of glucose and mannose induced the formation of symmetric and thick colonies (Figure S3A). The enhanced production of the extracellular matrix was evident in LGG colony cells grown with glucose demonstrating an increased production of the extracellular matrix compared with TSB alone (Figure S3B). Glucose and mannose also induced proteins related to adhesion and biofilm formation (25, 26) (Figure 33C).

To further confirm that the altered levels of cell wall biosynthesis proteins during fermentation reflect physiological changes in peptidoglycan composition, and to determine whether cell wall remodeling inducing biofilm formation or vice versa, we analyzed muramopeptides composition using LC-MS during both planktonic growth and biofilm formation. Our analysis demonstrated large distinction in cell composition during application of fermentable sugars, independently to the state of growth (Figure 5G, H). These results highlight the importance of cell wall remodeling during growth on fermentable sugars, and that it occurs prior to biofilm formation or bacterial aggregation.

### Specific metabolome-associated changes in the antimicrobial properties of LGG

To examine whether LGG secretes proteinaceous antibacterial substances together with organic acids during fermentation, we tested whether fermentation could alter the outcomes of microbial competition of LGG against enteric pathogens. LGG colonies grown with fermentable glucose in the medium are more competitive in the presence of gut pathogens. In the absence of glucose or application of non-fermentable raffinose, *S. typhimurium* colonies attached to and engulfed LGG colonies. Under these conditions, *E. faecalis* colonies were merging with and invading to colonies formed by LGG (Figure 6A). In contrast, in the presence of glucose, *S. typhimurium* colonies were strongly antagonized and *E. faecalis* by LGG during competition (Figure 6A) (Figure 6A). In addition, we tested the competitiveness of LGG during fermentation against additional bacteria that do not share the same niche, i.e., *P. aeruginosa*. Consistent with the increased broad-spectrum effects versus enteric pathogens, *P. aeruginosa* is clearly antagonized by LGG in a fermentation-dependent manner (Figure 6A).

**Figure 6.**
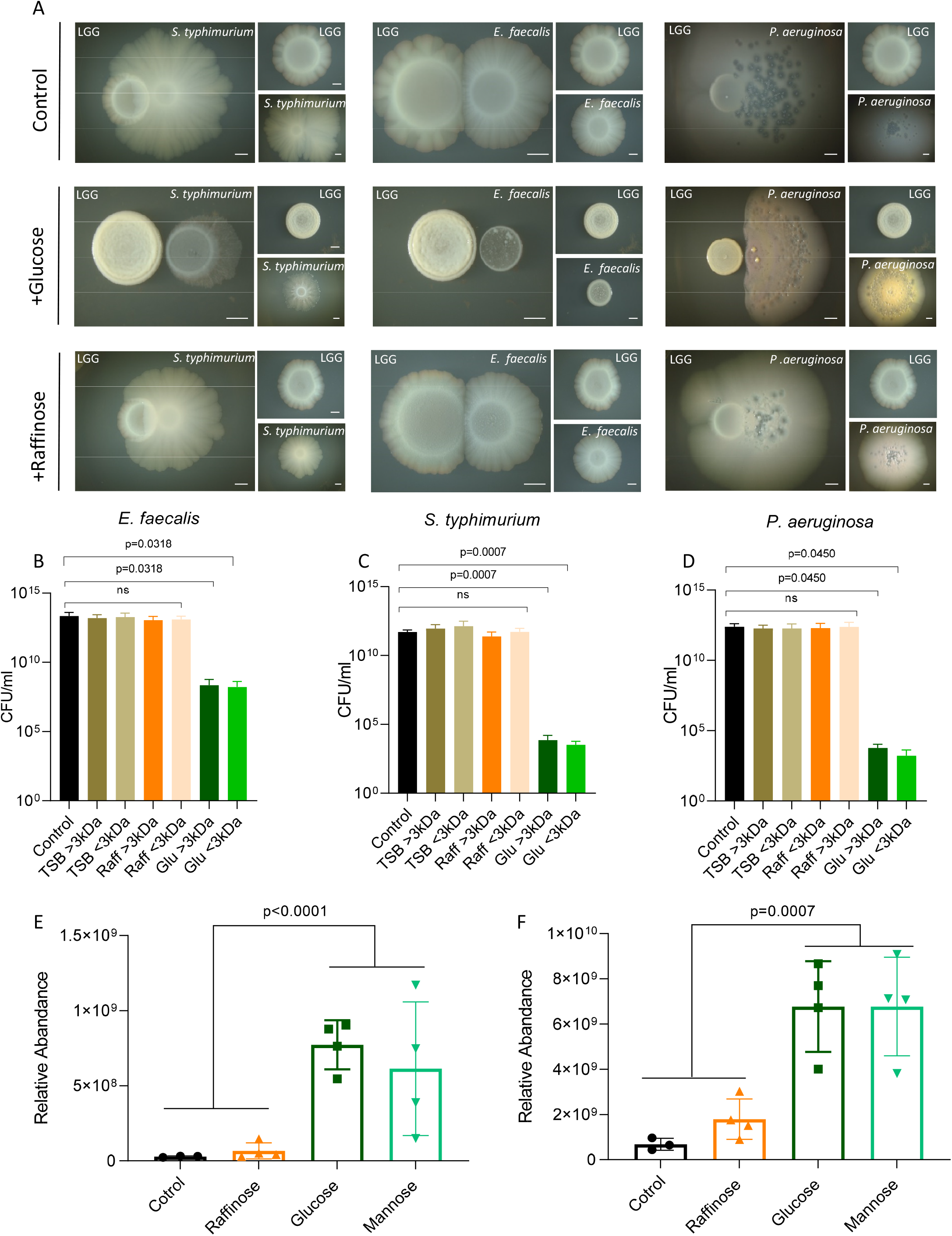
Fermentation has a cardinal role in antimicrobial production. **(A)** Colonies of LGG, *S. typhimurium, E. faecalis and P. aeruginosa*, grown on solid TSB medium or TSB supplemented with glucose (1% W/V) or raffinose (1% W/V). The colonies were incubated at 37°C, in in CO_2_ enriched environment, for 7 days. Colonies of LGG were inoculated next to *S. typhimurium E. faecalis* colonies at a distance of 0.2 cm. **(B-D)** Colony-forming units counts of *E. faecalis* or *S. typhimurium* or *P. aeruginosa* grown in 30% CM >3 kDa or <3kDa from LGG grown in TSB medium or TSB supplemented with glucose (1% W/V) or raffinose (1% W/V). Statistical analysis was performed using Brown-Forsythe and Welch’s ANOVA with Dunnett’s T3 multiple comparisons test. p < 0.05 was considered statistically significant. **(E)** protein relative abundance between sample of Msp1/p75 and **(F)** Msp2/p40. Statistical analysis was performed using Student’s t-Test followed by FDR correction. p< 0.05 was considered statistically significant.

The inhibitory product was secreted as predicted by the proteome as the conditioned medium of LGG with glucose (but not glucose) inhibited the growth of all three pathogens (Figure S4). Conditioned medium of LGG stationary cultures in the absence of fermentable sugar had little or no effect on the growth of these competitors. To confirm a fermentation-dependent proteinous component in the toxicity of LGG, we separated the condition medium by size of 3 KDa to separate between small molecules (primarily organic acids, and proteins). Fractions bigger and smaller than 3 kDa obtained from conditioned medium with glucose significantly inhibited the growth all three pathogens while fractions from conditioned media collected with no added sugar or raffinose had a significantly reduced activity (Figure 6B-D). The proteome analysis supported the notion of antibacterial protein effectors being induced during fermentation.

Specifically, Msp1/p75 (Figure 6E) and Msp2/p40 (Figure 6F) were induced by fermentation and Msp1/p75 was detected independently in the >3KDa fraction (Figure S5). Those proteins secreted by LGG (27) and are probiotic effectors (28), with peptidoglycan hydrolases activity (29). Msp1/P75 was characterized as a D-glutamyl-L-lysyl endopeptidase with a role in cell wall metabolism (30), has an inhibitory effect on *Candida albicans* by chitinase activity (31) and surface display of this protein in *Bacillus subtilis* enhanced its antibacterial activity against *Listeria monocytogenes*(32).

## Discussion

Diet emerges as a pivotal determinant of gut microbiota community structure and function (33). The influence of nutrients and other environmental factors that qualitatively change the composition of gut microbiota indicated that gut microbiota influences the metabolism of cells in the liver and adipose tissues (34). Gut microbiota contributes to host metabolism, including food intake, digestion, absorption and energy expenditure through the production of a myriad of metabolites, such as short-chain fatty acids, bile acids, amino acids and their derivatives (35, 36). It was found that microbiota-derived metabolites such as acetate, lactate, succinate and myristoleic acid can regulate intestinal absorption and lipid metabolism (37). However, how gut microbiota and its derived metabolites are altered by specific dietary changes is largely undetermined.

Probiotic bacteria are a subset of GI microbiota proposed to benefit human health. Here we use the probiotic bacterium LGG as our lens to the link between dietary changes, microbial physiology and microbial metabolites. LGG is one of the most widely used probiotic strains. Various health effects are well documented including the prevention and treatment of gastro-intestinal infections and diarrhea, and stimulation of immune responses that promote vaccination or even prevent certain allergic symptoms. However, not all interventions studies have demonstrated clinical benefit probiotics, and even for the same conditions, the results are not always consistent (28). Linking fermentable sugars with LGG performance in vitro we can generate specific predictions regarding specific dietary associated probiotic behavior. First, probiotics can clearly exclude or inhibit pathogens, either through direct action or through influence on the commensal microbiota (38–40).

We show here that the availability of fermentable sugar is a cardinal determinant to this behavior, and can be attributed to enhanced accumulation of cytotoxic metabolites (Ethanol, organic acids and butanol-Figure 2), volatiles (Figure 3) and extracellular proteins (Figure 6E,F). While a broad spectrum of pathogenic bacteria are eliminated by LGG in a fermentation-dependent manner (Figure 6), the bacterium itself alters its cell wall (Figure 5C,D,G,H) and biofilm formation (Figure 5E,F), as a potential coping mechanism against the same conditions. Furthermore, our results indicated that cell wall remodeling precedes biofilm formation, and that it is very likely that metabolic rewiring and the accumulation of toxic fermentation products (11) trigger the cell wall remodeling. Also supporting this notion, is that one of the antimicrobial proteins induced during fermentation (Figure 6C) is a D-glutamyl-L-lysyl endopeptidase, targeting peptidoglycan. Biofilm formation is tightly linked with cell wall remodeling and is expected to be co-regulated with antimicrobial activity (41, 42). Furthermore, acetate (accumulated intracellularly and emitted as volatile) activates the biosynthesis of the bacteriocin rhamnosin B in LGG when applied as acetic acid salt (43). Consistently, our data provide strong evidence that cell wall remodeling precedes an induced biofilm formation (Figure 5), and these behaviors are correlated with enhanced antimicrobial production (Figure 6).

A second mechanism by which LGG exerts host-beneficial effects is modulate host immune responses, exerting strain-specific local and systemic effects (44). Many of the interactions between probiotic bacteria and intestinal epithelial and immune cells are thought to be mediated by molecular structures, known as microbe-associated molecular patterns (MAMPs), which can be recognized through specific pattern recognition receptors (PRRs) such as Toll-like receptors (TLRs) (45–47). These behaviors greatly depend on the capacity of bacterium to adhere to host tissues manifested in enhanced biofilm formation, which occurs downstream to fermentation (Figure 5E, F) and the fermentation-dependent expression of proteins associated with an increased adhesion (Figure S3).

Experimental *in vitro* data and experiments in animal models validate diet-bacterial response-host interactions for probiotic strains, including LGG. However, the vast majority of published data are from *in vivo* studies in humans with poor deception for the bacterial mechanisms of action. We here show that for an optimized and more tailored application of probiotics, it is imperative to understand the mechanisms of interaction with the host and host’s diet in detail. In here we provide compelling evidence that linking specific dietary changes with the performance of probiotic strains. Our results may yield improved interventions based on probiotic bacteria, or their products.

## Materials and Methods

### Strains, media and imaging

*Lacticaseibacillus rhamnosus* GG (LGG) ATCC 53103 probiotic strain used in the study. *Enterococcus faecalis* ATCC 29212142, *Salmonella enterica* serovar Typhimurium [kindly provided by Dr. Roi Avraham’s Lab] and *Pseudomonas aeruginosa* PA14 were used for competition assay. A single colony of *L. rhamnosus* GG was isolated on a solid deMan, Rogosa, Sharpe (MRS) plate (1.5% agar), inoculated into 5 mL MRS broth (Difco, Le Pont de Claix, France), and grown at 37°C, without shaking overnight. A single colony of *E. faecalis, S. typhimurium* and *P. aeruginosa* isolated on a solid LB agar plate was inoculated into 3ml Luria-Bertani (LB) (Difco) and grown at 37°C, with shaking overnight. For biofilm colonies, these cultures were inoculated into a solid medium (1.5% agar) containing 50% Tryptic soy broth (TSB), TSB supplemented with (1% w/v) D-(+) - glucose, (1% w/v) D-(+)- raffinose or (1% w/v) D- (+)- mannose. The bacteria were incubated in a BD GasPak EZ - Incubation Container with BD GasPak EZ CO2 Container System Sachets (260679) (Becton, Sparks, MD, USA), for 72 hours or 7 days, in 37°C. The colony images were taken using a Stereo Discovery V20” microscope (Tochigi, Japan) with objectives Plan Apo S ×1.0 FWD 60 mm (Zeiss, Goettingen, Germany) attached to a high-resolution microscopy Axiocam camera. Data were created and processed using Axiovision suite software (Zeiss). For planktonic growth, the bacterial cultures were inoculated into a liquid medium 50% TSB with different sugars as described above, incubated for 24h, no shaking, at 37°C.

### Growth measurement and analysis

LGG cultured cells grown for overnight were diluted 1:100 in 200 μL liquid medium contains 50% TSB (BD), TSB supplemented with (1% w/v) different sugars in a 96-well microplate (Thermo Scientific, Roskilde, Denmark). *S. typhimurium, E. faecalis and P. aeruginosa* cultured cells grown for overnight were diluted 1:100 in 200 μL liquid medium contains TSB supplemented with (1% w/v) and with 30% conditioned medium derived from LGG grown with and without sugars as mentioned in the first section. Cells were grown with agitation at 37 °C for 18 h in a microplate reader (Tecan, Männedorf, Switzerland), and the optical density at 600 nm (OD_600_) was measured every 30 min.

### Fluorescence microscopy

A bacterial biofilm colony of LGG grown as described above was suspended in 200 μL 1x Phosphate-Buffered Saline (PBS), and dispersed by pipetting. Samples were centrifuged briefly, pelleted and re-suspended in 5μL of 1x PBS supplemented with the membrane stain FM1-43 (Molecular Probes, Eugene, OR, USA) at 1 μg/ml. The cells were then placed on a microscope slide and covered with a poly-L-Lysine (Sigma) treated coverslip. The cells were observed by Axio microscope (Zeiss, Germany). Images was analyzed by Zen-10 software (Zeiss).

### Proteomics - sample preparation, LC/MS and data analysis

LGG grown in TSB or TSB with 1% sugar for 24 h in 37°C. To get an equal number of bacteria the OD_600_ of the cultures was compared. The cell pellets were subjected to in-solution tryptic digestion using the suspension trapping (S-trap) method as previously described (48). Briefly, bacterial cell pellets were homogenized in the presence of lysis buffer containing 5% SDS in 50mM Tris-HCl, pH 7.4. Lysates were incubated at 96°C for 5 min, followed by six cycles of 30 s of sonication (Bioruptor Pico, Diagenode, USA). Protein concentration was measured using the BCA assay (Thermo Scientific, USA). 50 ug of total protein was reduced with 5 mM dithiothreitol and alkylated with 10 mM iodoacetamide in the dark. Each sample was loaded onto S-Trap microcolumns (Protifi,USA) according to the manufacturer’s instructions. After loading, samples were washed with 90:10% methanol/50 mM ammonium bicarbonate. Samples were then digested with trypsin (1:50 trypsin/protein) for 1.5 h at 47°C. The digested peptides were eluted using 50 mM ammonium bicarbonate. Trypsin was added to this fraction and incubated overnight at 37°C. Two more elutions were made using 0.2% formic acid and 0.2% formic acid in 50% acetonitrile. The three elutions were pooled together and vacuum-centrifuged to dryness. Samples were resuspended in H2O with 0.1% formic acid and subjected to solid phase extraction (Oasis HLB, Waters, Milford, MA, USA) according to manufacturer instructions and vacuum-centrifuged to dryness. Samples were kept at−80°C until further analysis.

### Liquid chromatography

ULC/MS grade solvents were used for all chromatographic steps. Dry digested samples were dissolved in 97:3% H2O/acetonitrile + 0.1% formic acid. Each sample was loaded using split-less nano-Ultra Performance Liquid Chromatography (10 kpsi nanoAcquity; Waters, Milford, MA, USA). The mobile phase was: A) H2O + 0.1% formic acid and B) acetonitrile + 0.1% formic acid. Desalting of the samples was performed online using a reversed-phase Symmetry C18 trapping column (180 µm internal diameter, 20 mm length, 5 µm particle size; Waters). The peptides were then separated using a self-packed analytic column containing ReproSil-Pur 120 C18-AQ resin (100 µm internal diameter column, 250 mm length, 1.9 µm particle size; Dr. Maisch, Germany) at 0.35 µL/min. Peptides were eluted from the column into the mass spectrometer using the following gradient: 4% to 30%B in 155 min, 30% to 90%B in 5 min, maintained at 90% for 5 min and then back to initial conditions.

### Mass Spectrometry

The nanoUPLC was coupled online through a nanoESI emitter (10 μm tip; New Objective; Woburn, MA, USA) to a quadrupole orbitrap mass spectrometer (Q Exactive HF, Thermo Scientific) using a FlexIon nanospray apparatus (Proxeon). Data was acquired in data dependent acquisition (DDA) mode, using a Top10 method. MS1 resolution was set to 120,000 (at 200m/z), mass range of 375-1650m/z, AGC of 3e6 and maximum injection time was set to 60msec. MS2 resolution was set to 15,000, quadrupole isolation 1.7m/z, AGC of 1e5, dynamic exclusion of 50sec and maximum injection time of 60msec.

### Data processing and analysis

Raw data was processed with MaxQuant v1.6.6.0 (49). The data was searched with the Andromeda search engine against LGG protein database from NCBI Reference Sequence: NC_017482.1 and appended with common lab protein contaminants. Enzyme specificity was set to trypsin and up to two missed cleavages were allowed. Fixed modification was set to carbamidomethylation of cysteines and variable modifications were set to oxidation of methionines, and deamidation of glutamines and asparagines. Peptide precursor ions were searched with a maximum mass deviation of 4.5 ppm and fragment ions with a maximum mass deviation of 20 ppm. Peptide and protein identifications were filtered at an FDR of 1% using the decoy database strategy (MaxQuant’s “Revert” module). The minimal peptide length was 7 amino-acids and the minimum Andromeda score for modified peptides was 40. Peptide identifications were propagated across samples using the match-between-runs option checked. Searches were performed with the label-free quantification option selected. The quantitative comparisons were calculated using Perseus v1.6.0.7. Decoy hits were filtered out and only proteins that had at least 2 valid values after logarithmic transformation in at least one experimental group were kept. Principle component analysis (PCA) of the log intensity values and heat map was performed with R. A Student’s t-Test followed by FDR correction, after the logarithmic transformation, was used to identify significant differences between the experimental groups, across the biological replica. Fold changes were calculated based on the ratio of geometric means of the different experimental groups. Go-term annotation of NC_017482.1 proteins was performed with Blast2Go tool. GO-terms over representation analysis (ORA) was performed with the WebGestalt (WEB-based Gen Set Analysis Toolkit) (14) on the up regulated set of proteins (FDR>0.05, fold change >2) and using a custom database of the genome’s GO annotation performed with Blast2Go tool. The cellular location for upregulated proteins (FDR>0.05, fold change >2) was predicted using PSORT Server v. 3.0 at https://www.psort.org/psortb/ (50).

### Peptidoglycan purification, preparation of muropeptides and LC-MS

Peptidoglycan (PG) sacculi were purified from planktonic cultures and biofilm colonies. Cells were collected by centrifugation (10,000 × g, 5 min) and kept in -20°C. frozen cells pellet washed with 1M NaCl, resuspended in 8% SDS in 0.1M Tris/HCl pH 6.8, boiled for 30 min. The suspension was then centrifuged (10,000 × g, 5 min) to collect the pellet, and washed five times with dH2O to remove SDS. Then the samples entered a sonifier water bath for 30 minutes and centrifuged. The pellet was then suspended in 15µg/mL DNAse, 60 µg/mL RNAse in 0.1M Tris/HCl pH 6.8 and incubated for 60 minutes at 37 °C, with gentle shaking. This was followed by treatment with 50 μg/mL and incubated at 37 °C for additional 60 minutes, with gentle shaking. To inactivate the enzymes the suspension boiled for 3 minutes, then centrifuged (5 min at 10,000 rpm) and washed once with dH2O. Samples were than boiled again in 4% for 30 min, and washed five times with dH2O to remove SDS. The PG pellet lyophilized, weighted and stored at -20 until papered for MS.

### Preparation for MS

The pellet resuspended in digestion buffer (12.5 mM sodium dihydrogen-phosphate, pH 5.5) with mutanolysin solution (5.000 U/ml) for 16h at 37°C with gentle shaking. To reduce muropeptides, equal volume of the solution of disaccharide peptides and of borate buffer (0.5 mM, pH 9.0) were mixed and incubated for 20 minutes at room temperature. pH adjusted to <4 with (1:5) phosphoric acid and filtered through 0.22µm.

### HPLC-MS/MS

An UltiMate 3000 UHPLC+ focused LC-MS system (Thermo Scientific™) coupled with a Q Exactive™ Focus Hybrid Quadrupole-Orbitrap™ Mass Spectrometer (Thermo Scientific™) was used for LC/HRMS analysis. Muropeptides were separated using a C18 analytical column (Accucore TM C18, 2.6µm particles, 100 × 2.1 mm; Thermo Fisher Scientific), column temperature at 50°C. The flow rate was 0.2 mL/min when solvent A is 100% water with 0.1% formic acid, and solvent B is 100% acetonitrile, and 0.1% formic acid. 10 µl sample injected, MS/MS data were acquired during a 40 min with gradient of: 0–12.5% B for 25 min, 12.5–20% B for 5 min, and stayed at 20% B for 5 min, and the column was re-equilibrated for 10 min under the initial conditions. The first 2 and last 5 min went to waste. The Q Exactive Focus was operated under positive ionization mode. The measurement was set to top 3 MS/MS fragmentations. The NCE was set with collision energy of 10,17.5,25. All MS spectra were analyzed by Xcalibur and freestyle software (Thermo Scientific™). PCA analysis created using MetaboAnalyst5.0.

### Transmission Electron Microscopy (TEM)

LGG biofilm colony was grown on solid medium (1.5% agar) contains 50% tryptic soy broth (TSB) with or without (1% w/v) D-(+) - glucose, for 7 days. Cells were fixed with 3% paraformaldehyde, 2% glutaraldehyde in 0.1 M cacodylate buffer containing 5 mM CaCl2 (pH 7.4), then post fixed in 1% osmium tetroxide supplemented with 0.5% potassium hexacyanoferrate tryhidrate and potassium dichromate in 0.1 M cacodylate (1 hour), stained with 2% uranyl acetate in water (1 hour), dehydrated in graded ethanol solutions and embedded in Agar 100 epoxy resin (Agar scientific Ltd., Stansted, UK). Ultrathin sections (70-90 nm) were viewed and photographed with a FEI Tecnai SPIRIT (FEI, Eidhoven, Netherlands) transmission electron microscope operated at 120 kV and equipped with an EAGLE CCD Camera.

### Image analysis for cell wall thickness quantification

We quantified the average and standard deviation of bacteria cell wall thickness from TEM images, by manually drawing the inner and outer cell wall borders, and then automatically matching the pairs of inner and outer boundaries to create a ring-like region of interest and quantifying the local thickness along the centerline (skeleton) of each cell wall. Manual drawing was done in Fiji (51) and the boundaries were saved as region of interests (ROIs) file. Automatic quantification was done using Fiji macro that read the TEM image together with the matching ROIs file. Color-coded visualization of the results is created using MorphoLibJ (52) plugin. The macro is available at: https://github.com/WIS-MICC-CellObservatory/BacteriaCellWallThickness. All the images were manually calibrated using to the scale bar, in order to get the measurements with proper calibration.

### Imaging Flow Cytometry

Biofilms colonies were cultures and incubated as mentioned in the first section. Colonies were harvested after 72 hours and separated with mild sonication. Colonies were harvested after 72 hours and separated with mild sonication. For cell wall labelling, cells were gently centrifuged, resuspended in 100 µl of PBS supplemented with WGA-FITC (5 µg/ml, Sigma), incubated for 15 min at room temperature, and washed twice with PBS before imaging. Data were acquired by ImageStreamX Mark II (AMNIS, Austin, Tx) using a 60× lens (NA=0.9). The laser used was at 785 nm (5 mW) for side scatter measurement. During acquisition, bacterial cells were gated according to their area (in square microns) and side scatter, which excluded the calibration beads (that run in the instrument along with the sample). For each sample, 100,000 events were collected. Data were analyzed using IDEAS 6.2 (AMNIS). Focused events were selected by the Gradient RMS, a measurement of image contrast. Cells stained with WGA-FITCH were selected using the Intensity (the sum of the background subtracted pixel values within the image) and Max Pixel values (the largest value of the background-subtracted pixels) of the green channel (Ch02). Cell wall intensity was quantified using the Mean Pixel feature (the mean of the background-subtracted pixels contained in the input mask).

### Biofilm formation assay

For biofilm growth, 1 µL of LGG starter culture was diluted (1:100) in 200 µL 50% TSB, TSB supplemented with (1% w/v) D-(+) - glucose, (1% w/v) D-(+)- raffinose or (1% w/v) D- (+)- mannose in 96-well polystyrene plates and incubated for 72 h in 37°C. Biofilm formation was assessed by crystal violet staining. After 72h planktonic cells were removed by pipetting, and wells washed with DDW (Deuterium-depleted water). The adherent cells were stained with 0.05% crystal violet stain for 30 min. The stain removed, and the wells were washed with DDW. 100% ethanol was added to the wells for 15 minutes. Crystal violet intensity was determined by a spectrophotometer (OD 575nm). For calculating the ratio between planktonic cells and attached cells, 2 µL of LGG starter culture was diluted (1:100) in 2 mL of 50% TSB, TSB supplemented with (1% w/v) D-(+) - glucose, (1% w/v) D-(+)- raffinose or (1% w/v) D- (+)- mannose in 12-well polystyrene plates and incubated for 24 h in 37°C. The plektonic cells were removed and their OD_600_ measured. The adherent cells resuspended in PBS (2mL) and their OD_600_ measured. The ratio calculated was adherent cells OD_600_/Planktonic cells OD_600_.

### Interaction assay

LGG, *S. typhimurium, E. faecalis and P. aeruginosa* biofilms colonies grown as mentioned in the first section. These bacteria were inoculated solid medium (1.5% agar) contains 50% tryptic soy broth (TSB) with or without (1% w/v) D- (+) - glucose or D-(+)- raffinose next to each other at the distance of 0.2 mm for 7 days. As a control each bacterium inoculated separately. All images were taken Stereo Discovery V20” microscope (Tochigi, Japan) with objectives Plan Apo S ×0.5 FWD 134 mm or Apo S ×1.0 FWD 60 mm (Zeiss, Goettingen, Germany) attached to a high-resolution microscopy Axiocam camera. Data were created and processed using Axiovision suite software (Zeiss).

### Conditioned media (CM) acquisition

LGG planktonic cultures, incubated for 24h, at 37°C were spun for 20 min at 4 °C at 4000 g to remove cells. The supernatant was than filtered through a 0.22 µm (Corning incorporated, USA). Then the supernatant was separated by size to a large (>3 kDa) and small (<3 kDa) fraction using Amicon ultrafiltration system with 3 kDa filter (Millipore, Ireland). The fraction then filtered again through a 0.22 µm (Millipore).

### CFU assay

*S. typhimurium, E. faecalis and P. aeruginosa* bacterial cultures were inoculated into a liquid medium 50% TSB supplemented with (1% w/v) D-(+) – glucose with 30% conditioned media (CM), incubated for 24h, no shaking, at 37°C. Then, the samples were serially diluted x10 into 96 well plates and 20 µL from each sample was plated on solid LB agar (1.5 % agar) using a multichannel pipette with the dot-spot technique. CFU enumeration was carried out following overnight incubation at 37°C.

### LC-MS for polar metabolite analysis

LGG planktonic cultures grown in 50% TSB, TSB supplemented with (1% w/v) D-(+) - glucose, (1% w/v) D-(+)- raffinose or (1% w/v) D- (+)- mannose for 24h at 37°C. To get an equal number of bacteria the OD_600_ of the bacteria was compared, and after that washed twice with PBS. For polar metabolite analysis in the polar phase samples, the lyophilized pellets were dissolved using 100 µL DDW-methanol (1:1), centrifuged twice (at maximum speed) to remove possible precipitants, and were injected into LC-MS system. Polar analysis in the polar phase was done as described in Zheng et.al. (2015) with minor modifications described below. Briefly, analysis was performed using Waters Acquity I class UPLC System combined with a mass spectrometer (Thermo Exactive Plus Orbitrap) operated in a negative ionization mode. The LC separation was done using the SeQuant Zic-pHilic (150 mm × 2.1 mm) with the SeQuant guard column (20 mm × 2.1 mm) (Merck). The Mobile phase B: acetonitrile and Mobile phase A: 20 mM ammonium carbonate with 0.1% ammonia hydroxide in water: acetonitrile (80:20, v/v). The flow rate was kept at 200 μL*min−1 and gradient as follow: 0-2min 75% of B, 17 min 12.5% of B, 17.1 min 25% of B, 19 min 25% of B, 19.1 min 75% of B, 23 min 75% of B.

### Polar metabolites data analysis

The data processing was done using TraceFinder Thermo Fisher software. Polar metabolites were identified by accurate mass, retention time, isotope pattern, and verified using an in-house mass spectra library.

### Volatiles collection and analysis

The samples were prepared for analysis in 20mL headspace vialsThe headspace (300 mL) above cultures was actively sampled. Volatile compounds (VCs) analysis was conducted on a thermal desorption-gas chromatography time-of-flight mass spectrometer (GC-TOF-MS) platform (Leco BT, Germany) combined with Gerstel MPS autosampler (Germany).. The VOCs (volatile organic compounds) were collected using SPME PDMS/DVB (pink) fiber at 30 0C for 15 min and desorbed for 3 min using temperature 220 0C. The GC column (ZB-624PLUS column, 30 m, 0.32 mm internal diameter, 1.8 μm film thickness, Phenomenex) was held at an initial temperature of 40°C for 3 min, ramped to 205 °C at 7°C min and hold 205 °C for 1 min. The GC runtime was 28 min. The TOF-MS was in electron ionization mode set at 70 eV. The source temperature was set to 250 °C, and spectra were acquired in dynamic range extension mode at 10 scans s−1 over a range of 39–500 m/z.

### GC data processing

GC-TOF-MS data were acquired and analysed using ChromaTof (Leco, Germany). Chromatographic peaks and mass spectra were cross-referenced with National Institute of Standards and Technology (NIST 17) and Wiley libraries for putative identification purposes (matching factor >750 match).

### Statistical analysis

Statistical analyses were performed with GraphPad Prism 9.0 (GraphPad 234 Software, Inc., San Diego, CA). Relevant statistical tests are mentioned in the indicated legends of the figures.

