## Supporting Information for "Metabolic rewiring of the probiotic bacterium *Lacticaseibacillus rhamnosus* GG contributes to cell-wall remodeling and antimicrobials production"

### Supporting Figures (S1-S5)

### Supporting materials and methods

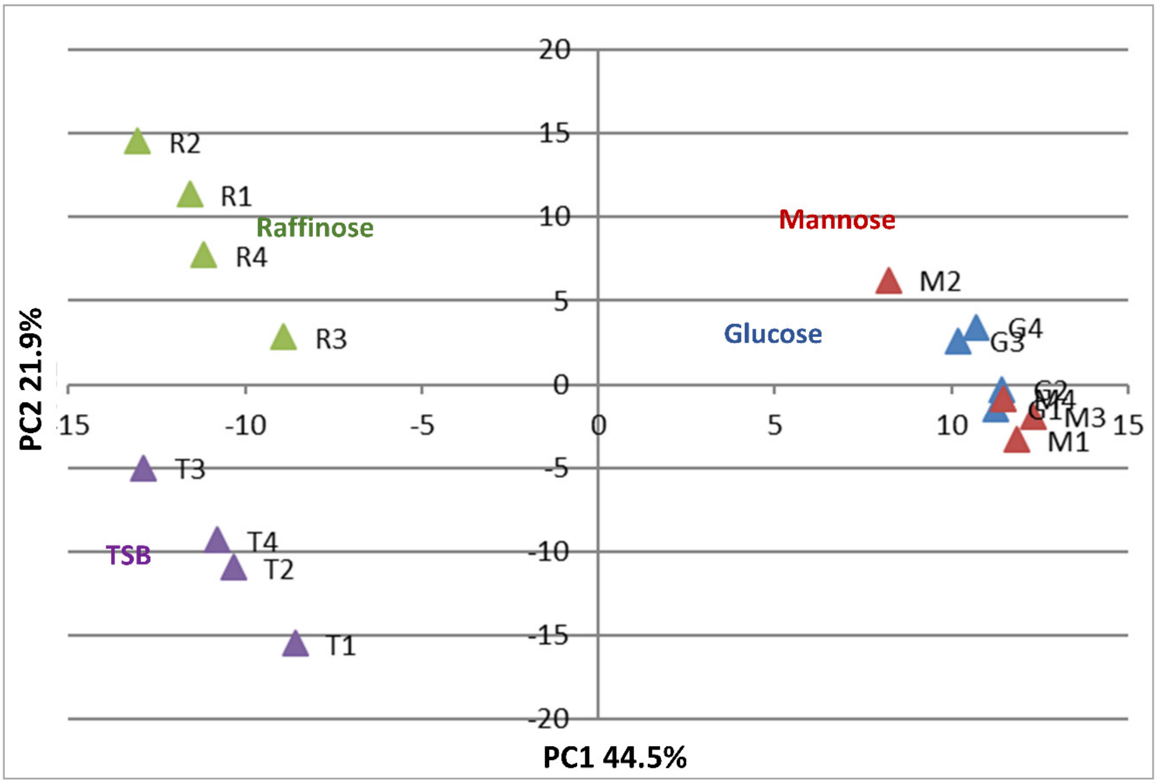


**Figure S1.** Principal component analysis plot of metabolomics analysis from LGG grown in liquid TSB medium or TSB medium supplemented with glucose (1% W/V), mannose (1% W/V) or raffinose (1% W/V). Raffinose [R1-R4], Glucose [G1-G4], Mannose [M1-M4], Non-supplemented TSB [T1-T4]

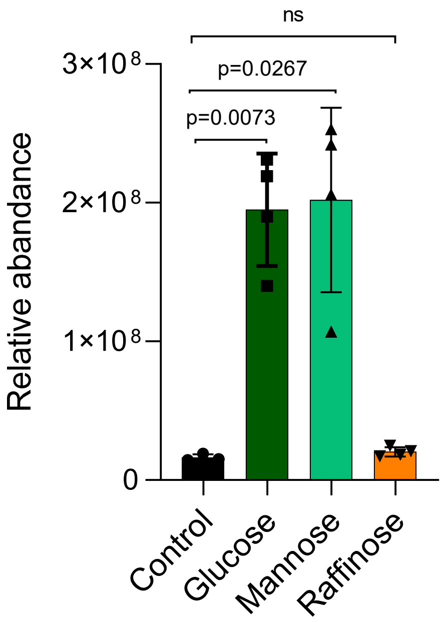


**Figure S2.** Protein relative abundance of Class A sortase between samples. Statistical analysis was performed using Brown-Forsythe and Welch's ANOVA with Dunnett's T3 multiple comparisons test. $p < 0.05$ was considered statistically significant.

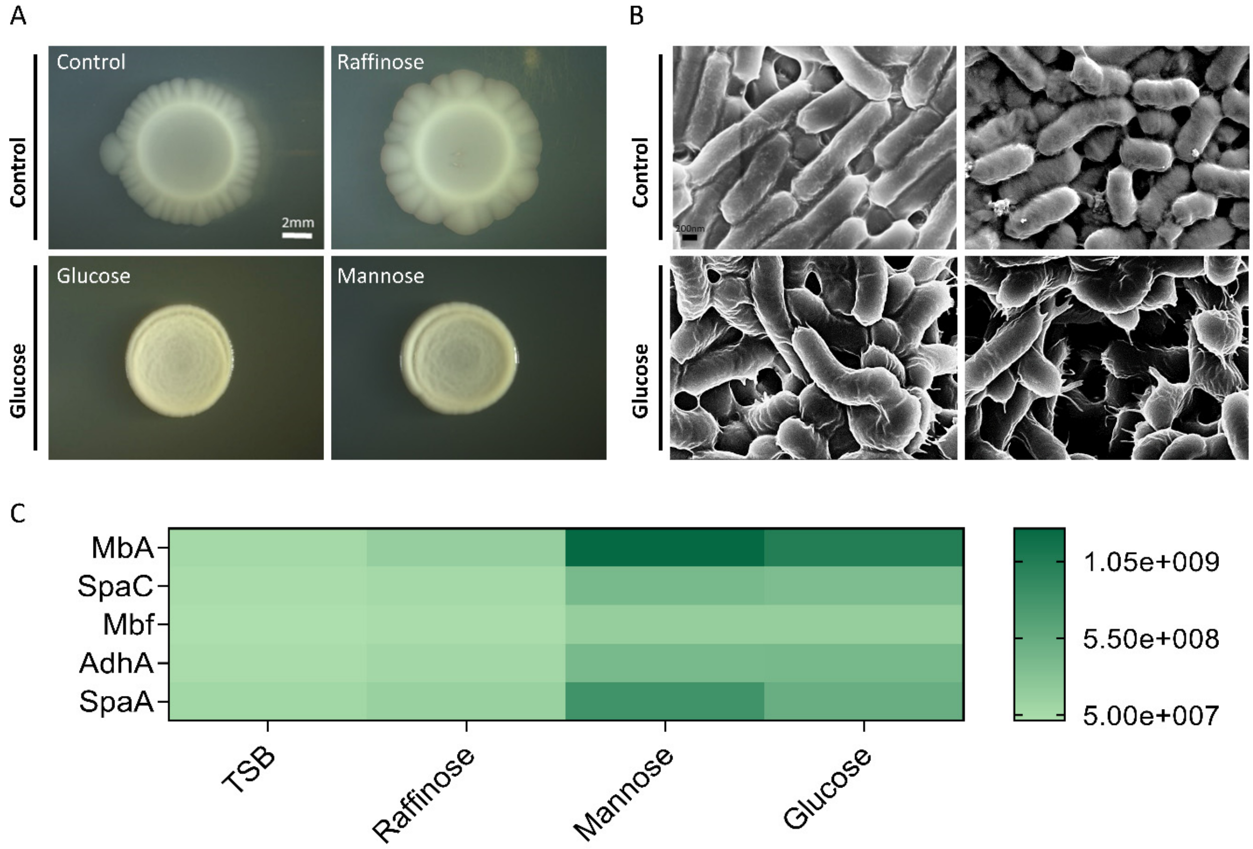


**Figure S3. (A)** LGG grown on solid TSB (control), TSB supplemented with glucose (1% W/V), mannose (1% W/V) or raffinose (1% W/V). Biofilms were grown at 37° C in CO2 enriched environment for 7 days. **(B)** Scanning electron microscopy (SEM) images of LGG grown in solid TSB (control), TSB supplemented with glucose (1% W/V). The colonies were incubated at 37°C in an environment enriched with $CO_2$ for 7 days. **(C)** Heat map based on fold differences in the intensity of adhesion proteins identified in proteomics analysis.

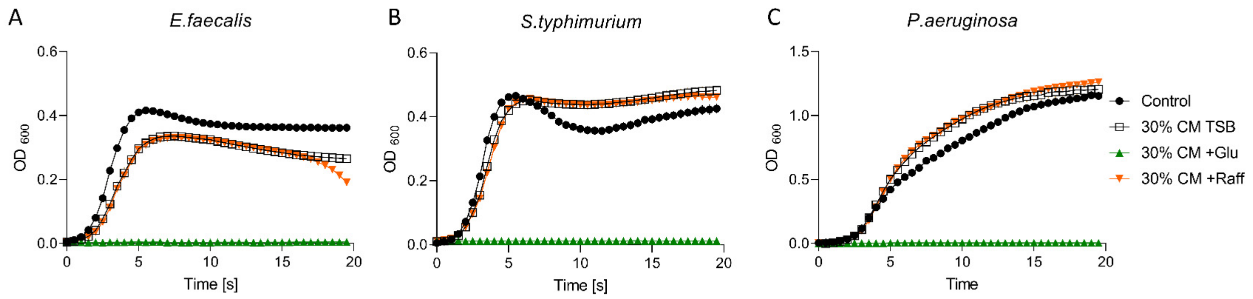


**Figure S4.** Growth curves of **(A)** *E. faecalis* , **(B)** *S. typhimurium* and **(C)** *P. aeruginosa* in 96 well plates in 37°C with shaking in TSB with glucose. Cells were supplemented with 30% conditioned medium (CM) derived from LGG grown in liquid TSB medium or TSB medium supplemented with glucose (1% W/V ) or raffinose (1% W/V).

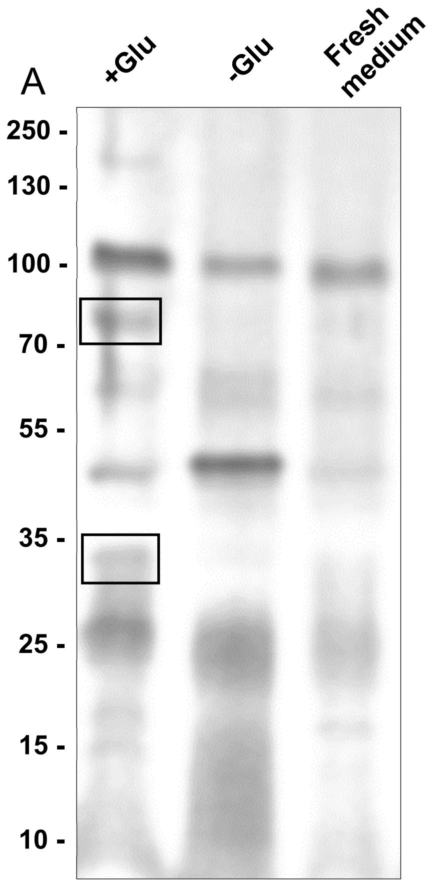


B

```
1   MVDSKKVLSV TAGFVGAAGL AALATGANTV SASTGTVSYK SGATTVWNSP SWHQVKRYVT
61   FGDTVQLLGK TVDQNGATWY KVGDNQWIPE LYLNVAGKTA TVETPSSAAS QTAVSQA
121 QAPTSQAPAT QTPAAPQTDT QTANTQLYVK NIGSAVTVWT TPAYTHATGQ YLEGSQTLTA
181 VAQQQANGET WYRLANGGYV PARFVSTTPV AVTPQPAAPQ SNEASVASTN TNAANDSAAA
241 SSAAASQAAA SSAAASTAAA NAAVASANAT ASQAAASEAA ASQAAASQAA ASQAAASQAA
301 ASQAAASQAA ASQAAASQAA ASQAAASQAA ANAAQQAPAN QANVTTTQVN ANQAQQQTAT
361 ATPAVNTSNQ TAAVSASRQA KIQAVIAIAE QQVGKPYVWG GKGPNSFDCS GLMYYAFLNG
421 AGVNIGGWTV PQESSGTQVS LSALQPGDLL FWGSHGSTYH VALYIGGGTM IQAPQPGENV
481 KYTALAYFMP DFAVRPSL
```

**Figure S5. (A)**SDS-PAGE of CM >3 kDa from LGG grown in TSB medium or TSB supplemented with glucose (1% W/V) and fresh TSB as a background. (B) Coverage of the identification of Msp1/P75 in the upper band.

### Supporting materials and methods

#### Scanning Electron Microscopy (SEM)

LGG biofilm colony was grown on solid medium (1.5% agar) contains 50% tryptic soy broth (TSB) with or without (1% w/v) D-(+) - glucose, for 3 days. Than the biofilms were fixed overnight at 4°C with 2% glutaraldehyde, 3% paraformaldehyde, 0.1 M sodium cacodylate (pH 7.4) and 5 mM CaCl2, dehydrated and dried as described by Bucher et al. 2015 (1). Mounted samples were coated with 3 nm thick Ir layer (Safematick). The imaging by secondary electron (SE) detector was preformed using a high-resolution Carl Zeiss Ultra 55 or Sigma scanning electron microscopes.

#### In gel proteolysis and mass spectrometry analysis

The big fraction driven from Lactobacillus rhamnosus GG grown with or without glucose and fresh 25% TSB+1% glucose as a control were mixed with 5x SDS sample buffer (10% SDS, 250 mM Tris-HCl pH 6.8, 0.5 M DTT, 0.1% bromophenol blue, 50% glycerol). Lysis and denaturation was carried out by incubation at 95 °C for 5 min. Proteins from the resulting cell extracts were loaded on a 10% SDS-PAGE gel, separated in SDS-running buffer (25 mM Tris, 192 mM glycine, 0.1% SDS) for 45 min with 80-120 V with a system from Bio-Rad. Proteins were stained with the Silver Stain. Silver-stained gel was destained with 30mM potassium hexacyanoferrate and 100mM sodium thiosulfate. The proteins in the gel were reduced with 2.8mM DTT (60ºC for 30 min), modified with 8.8mM iodoacetamide in 100mM ammonium bicarbonate (in the dark, room temperature for 30 min) and digested in 10% Acetonitrile and 10mM ammonium bicarbonate with modified trypsin (Promega) overnight at 37oC.

The resulting tryptic peptides were desalted using C18 tips (Homemade stage tips) dried and re-suspended in 0.1% Formic acid. The resulting tryptic peptides were resolved by reverse-phase chromatography on 0.075 X 200-mm fused silica capillary (J&W) packed with Reprosil reversed phase material (Dr Maisch GmbH, Germany). The peptides were eluted with linear 30 minutes gradient of 5% to 28% acetonitrile with 0.1% formic acid in water ,15 minutes gradient of 28% to 95% acetonitrile with 0.1% formic acid in water and 15 minutes at 95% acetonitrile with 0.1% formic acid in water at flow rates of 0.15 μl/min. Mass spectrometry was performed by a Q-Exactive plus mass spectrometer (Thermo) in a positive mode using repetitively full MS scan followed by High energy Collision Dissociation (HCD) of the 10 most dominant ion selected from the first MS scan. The mass spectrometry data was analyzed using Proteome Discoverer 1.4 software Using Sequest (Thermo) algorithm searching against the human proteome from the Uniport database, and L. rhamnosus GG from the NCBI-nr database. Semi quantitation was done by calculating the peak area of each peptide based its extracted ion currents (XICs). The area of the protein is the average of the three most intense peptides from each protein. Results were filtered with 1% false discovery rate.
